# CovRadar: Continuously tracking and filtering SARS-CoV-2 mutations for molecular surveillance

**DOI:** 10.1101/2021.02.03.429146

**Authors:** Alice Wittig, Fábio Miranda, Martin Hölzer, Tom Altenburg, Jakub M. Bartoszewicz, Sebastian Beyvers, Marius A. Dieckmann, Ulrich Genske, Sven H. Giese, Melania Nowicka, Hugues Richard, Henning Schiebenhoefer, Anna-Juliane Schmachtenberg, Paul Sieben, Ming Tang, Julius Tembrockhaus, Bernhard Y. Renard, Stephan Fuchs

**Author notes:** These authors contributed equally to this work (Alice Wittig, Fábio Miranda).

## Abstract

The SARS-CoV-2 pandemic underlined the importance of molecular surveillance to track the evolution of the virus and inform public health interventions. Fast analysis, easy visualization and convenient filtering of the latest virus sequences are essential for this purpose. However, access to computational resources, the lack of bioinformatics expertise, and the sheer volume of sequences in public databases complicate surveillance efforts. CovRadar combines an analytical pipeline and a web application designed for the molecular surveillance of the spike gene of SARS-CoV-2, an important vaccine target. The intuitive web front-end focuses on mutations rather than viral lineages and provides easy access to frequencies and spatio-temporal distributions from global sample collections. The data is regularly updated based on a scalable and reproducible analytical back-end. With this platform, we aim to give users, those with or without bioinformatics skills or sufficient computational resources, the possibility to track and explore mutational changes in the SARS-CoV-2 spike gene and to filter, download, and further analyze data that meet their questions and needs. Advanced computational users have the ability to apply the analytical pipeline and data visualization methods locally on their own data. CovRadar is freely accessible at https://covradar.net, source code is available at https://gitlab.com/dacs-hpi/covradar.

**GRAPHICAL ABSTRACT:** 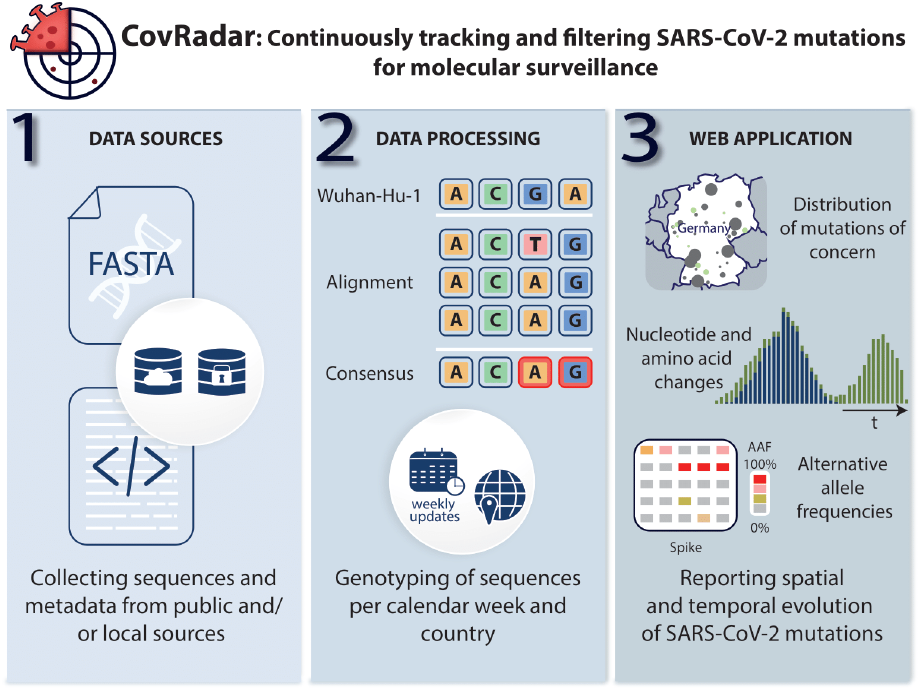

## INTRODUCTION

COVID-19 is an infectious disease caused by the novel coronavirus SARS-CoV-2, which was declared a global pandemic by the World Health Organization in March 2020. Two years later, more than 434 million cases and about 5.9 million deaths have been reported worldwide (WHO, Weekly operational update on COVID-19, March 1 2022).

Molecular surveillance plays a crucial role to understand evolutionary changes and answers epidemiological questions [1]. With decreasing costs and broader access to sequencing technologies, whole-genome sequencing has proven to be an indispensable tool for molecular surveillance of SARS-CoV-2 during the pandemic [2].

However, worldwide molecular surveillance efforts also revealed a bioinformatics bottleneck to rapidly develop specialized algorithms and tools to handle and analyze SARS-CoV-2 sequencing data and track changes in the pandemic faster [3].

In March 2022, the COVID-19 Data Portal hosted by EMBL-EBI [4] includes more than 4 million genome sequences and the international SARS-CoV-2 database GISAID [5, 6, 7] more than 9 million genome sequences, resulting in a 94 GB and a 250 GB multi-FASTA file, respectively. The sheer amount of data makes the analysis and search for known, new, and emerging mutations in the viral genome even more complicated and requires not only bioinformatics expertise but also significant computational resources; at the same time, comprehensive analysis of these data is essential to detect new viral variants, determine fixation rates, and ensure the efficacy of vaccines and diagnostic tests. Various tools were therefore developed in a short amount of time to tackle such challenges and to process the data in different ways while maintaining easy access to those with no computational background.

We performed a literature search and found 17 online tools available in the field of molecular surveillance of SARS-CoV-2 (Supplementary Table S1). For example, Nextstrain (nextstrain.org) [8] provides a platform for real-time tracking of SARS-CoV-2 evolution but needs to significantly down-sample sequences for calculations and visualizations. Other commonly used platforms such as outbreak.info and covariants.org also provide insights into mutations but group them based on already defined SARS-CoV-2 lineages [9] or lack options to examine detailed spatial distributions when the metadata contains such information. With Cov-Spektrum (https://cov-spectrum.org) [10], it is possible to monitor variants and mutations, but the tool lacks options to directly compare different countries, time periods, or hosts. We continue to see an urgent need to make SARS-CoV-2 sequence information more readily available to virologists and epidemiologists on a daily basis, especially in a public health context.

Here we present CovRadar, which was developed to address the aforementioned challenges. CovRadar consists of an analytical pipeline and an intuitive web application hosted on a powerful and scalable Kubernetes cluster (Fig. 1). The analytical workflow can process millions of genome sequences and can also combine different data sources. CovRadar functions primarily at the mutational level, allowing for early observations even before a new SARS-CoV-2 variant is assigned an official lineage name. Researchers can explore vast amounts of already pre-processed genome data, select specific subsets in space and time, and export their analyses for further investigations. Both, the pipeline and the web application can be installed and run on a local system to include access-restricted sequences and additional metadata. The data sources underlying CovRadar’s visualizations are updated regularly and therefore allow tracking of SARS-CoV-2 mutations in the spike gene in near real-time, so that threatening changes can be detected quickly and local health authorities can be informed and respond.

**Figure 1.**
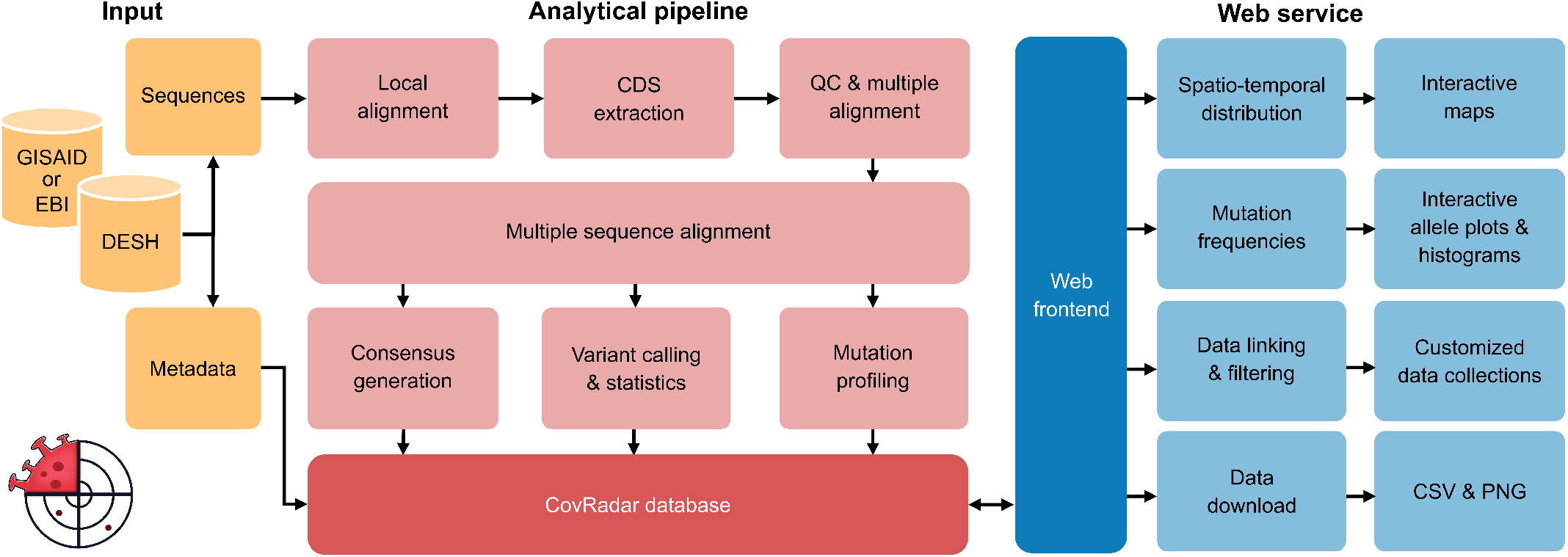
Simplified overview of Covradar’s input, analytical pipeline, and web service. The analytical pipeline is described in detail in the Supplementary Methods (Fig. S1). DESH – The German Electronic Sequence Data Hub which we combine with data from e.g. EMBL-EBI’s COVID-19 Data Hub for higher national resolution.

## MATERIALS AND METHODS

### Overview

The web application (covradar.net) is implemented using a Python-based web framework, Flask (github. com/pallets/flask), that comprises two Dash applications (https://github.com/plotly/dash), each connected to a MariaDB database. One database includes the results of the analytical pipeline (Fig. 1 and Fig. S1) for sequences of the COVID-19 Data Portal hosted by EMBL-EBI (covid19dataportal.org) and is updated bi-weekly, the other database includes the sequences of the last four weeks and is updated every other day. All back-end functionalities of the web application are written in Python. Images and tables are produced via Plotly.js (https://plot.ly). By having both the back- and front-end written in Python, we can more easily maintain the code and integrate further interactive user interfaces while keeping the Python syntax for the back-end logic and algorithms, instead of using PHP or JavaScript in the front-end. The design principles of CovRadar’s web application are focused on intuitive overviews that allow users to customize the visualized data based on their needs and to further download the filtered data sets for *post-hoc* specialized analyses. All elements such as plots and tables are interactive and can be directly modified and downloaded in PNG format and raw data format (CSV) for further processing. The web application runs on a Kubernetes cluster hosted by de.NBI. A local version can be started with the integrated Flask server or with a provided Docker container. Detailed instructions for installation can be found on https://gitlab.com/dacs-hpi/covradar.

### Position converter

To ease coordinate based queries, we implemented a converter between genome and protein sequences (accessible from the right sidebar of the website). The user can easily convert a range of genomic positions into (codon) positions within the spike gene or amino acids multiple sequence alignment (MSA) positions. For example, the amino acid change D614G corresponds to nucleotide positions 1840*−*1842 in the spike gene, which represents genomic coordinates 23402*−*23404 on the reference (Wuhan-Hu-1). The converter script can be also used as a stand-alone tool supporting all genes encoded by the SARS-CoV-2 genome (gitlab.com/dacs-hpi/covradar/-/tree/master/shared/residue_converter).

### Geographic distribution map of mutation frequencies

An interactive map shows the presence and frequency of selected amino acid changes for a given time period. For the geographic distribution, we use the highest resolution metadata available, which are postal codes for the German data (obtained via DESH, the German Electronic Sequence Data Hub) and countries otherwise (see Supplementary Methods). Importantly, the sampling locations for the German data are based on the postal codes of the primary diagnostic laboratories where a PCR test was performed to select a sample for sequencing. Therefore, the mutation frequencies shown for the Germany map do not necessarily correspond with the real distribution of SARS-CoV-2 mutations in Germany and do not allow direct conclusions about the mutation distribution in an area. Nevertheless, by integrating our national data source, we achieve a higher geographic resolution in comparison to what is possible only based on GISAID or EMBL-EBI data and for other countries. The user can select between multiple visualization modes. The mode ”n-th most frequent mutations per location” shows for n=1 the most frequent mutation per location, for n=2 the second most frequent mutation, and so on. The map can also show the frequencies of selected mutations per location. The “mutation proportion” mode can be used to check how many sequences have at least one of the selected mutations. Finally, the user can also examine the increase per location determined by linear regression within the selected time interval.

Clicking on a location within the map further shows information for that location in two plots: 1) A bar chart showing the number of sequences (mode “frequency”) and slopes (mode “increase”), respectively, as well as 2) the time course of mutation frequencies of a selected mutation and for a certain geographic location using linear regression.

### Alternative allele frequency plot

We implemented a heatmap-inspired alternative allele frequency (AAF) plot to visualize the frequencies of mutated sites (substitutions, INDELs) in the spike gene (Fig. 2). Each block represents a nucleotide in the MSA of extracted spike sequences. The coordinates on the y-axis correspond to the positions in the MSA. The colors represent the AAF of the selected sequences in comparison to a reference sequence (default: Wuhan-Hu-1, NC 045512.2). The mouse-over tooltips show the user the position references as well as the AAF and the coverage. Coverage describes how many sequences are used to retrieve the AAF at this position. Importantly, the analysis of the consensus sequences is not limited to the Wuhan-Hu-1 reference spike but it can also be compared to a large selection of consensus sequences that are provided per calendar week and country (see also Fig. S2). By that, the user can, for example, compare the current mutational pattern in Germany against the consensus sequence from another country in the same (or different) time period. The filters allow only specific SARS-CoV-2 lineages, hosts, time intervals, and countries to be selected. For better interpretation of the selected data, the number of sequences in the consensus sequence and in the filtered data set are shown above the plot. By default, the region encoding the receptor binding domain (RBD) of the spike gene is highlighted, but additional domains can also be activated, as well as any mutation positions shown in the geographic distribution map and custom amino acid/codon positions. This can be used to highlight specific locations when the plot is exported as a PNG. Further details about how AAF are calculated can be found in the Supplementary Methods.

**Figure 2.**
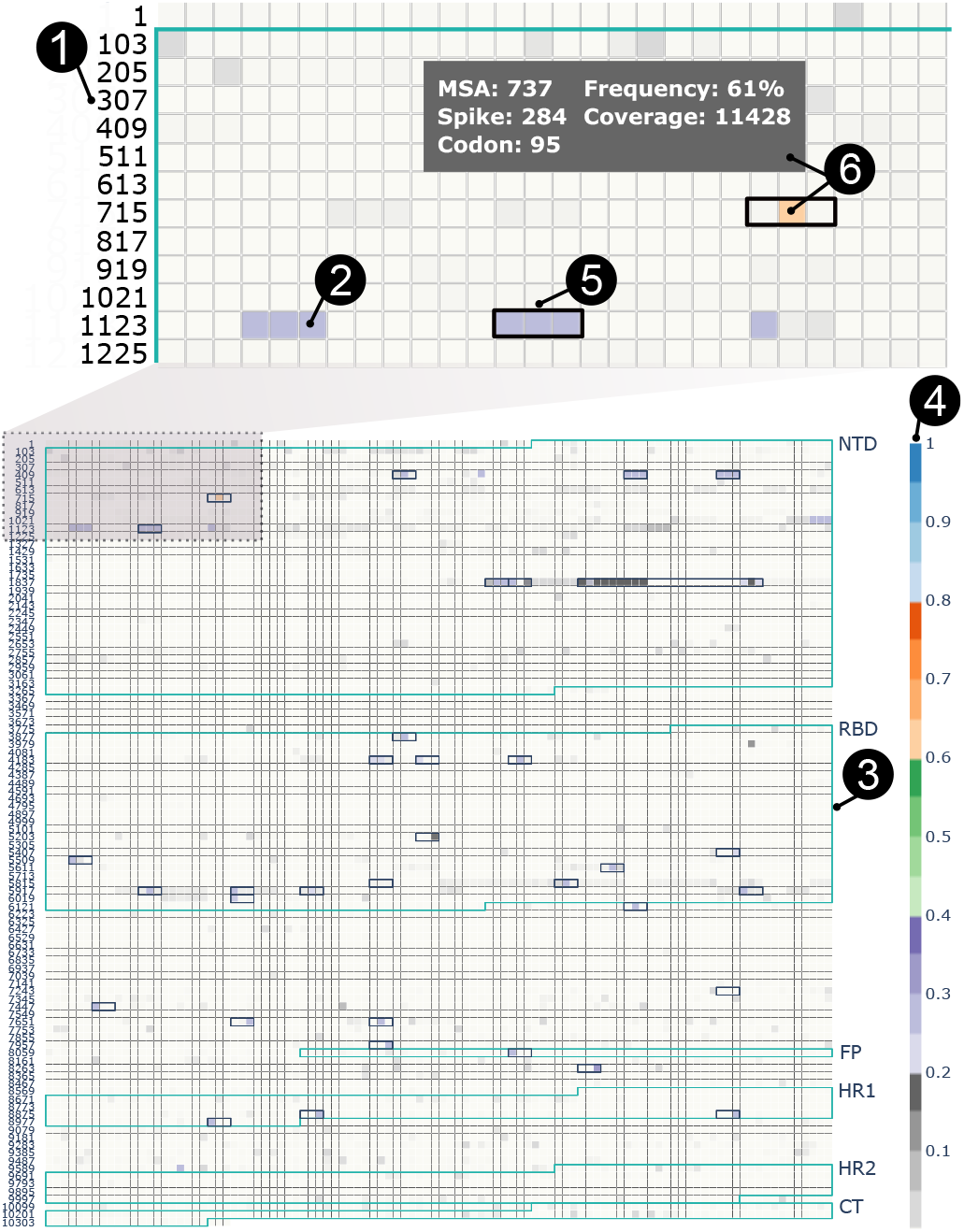
Alternative allele frequency (AAF) plot of German sequences from calendar weeks (CW) 51 and 52 of 2021 compared to the German consensus sequence of CW50. Coordinates on the left correspond to the MSA positions **(1)** and can be easily converted via the position converter or via mouse over. Each block represents a nucleotide position in the MSA of spike gene sequences **(2)**. Spike domains are highlighted with blue frames **(3)** and labeled NTD: N-terminal Domain, RBD: Receptor Binding Domain, FP: fusion peptide, HR1 and 2: heptapeptide repeat sequence 1 and 2, CT: cytoplasmic domain fusion. The color of the block represents the AAF **(4)**. The plot shows increased frequencies for nucleotide positions common for Omicron (mutations with *>* 75 % frequency based on outbreak.info, black frames **(5)**). The highest AAF in this comparison (∼61 %) belongs to codon 95 (orange box, **(6)**). Shown data is based on the DESH data set.

### Mutation distribution plots

The nucleotide distribution plot shows the frequency of different nucleotides at a given spike gene for each calendar week and country. Accordingly, an amino acid distribution plot highlight the frequency of different amino acids encoded by the respective codon (Fig. 3).

**Figure 3.**
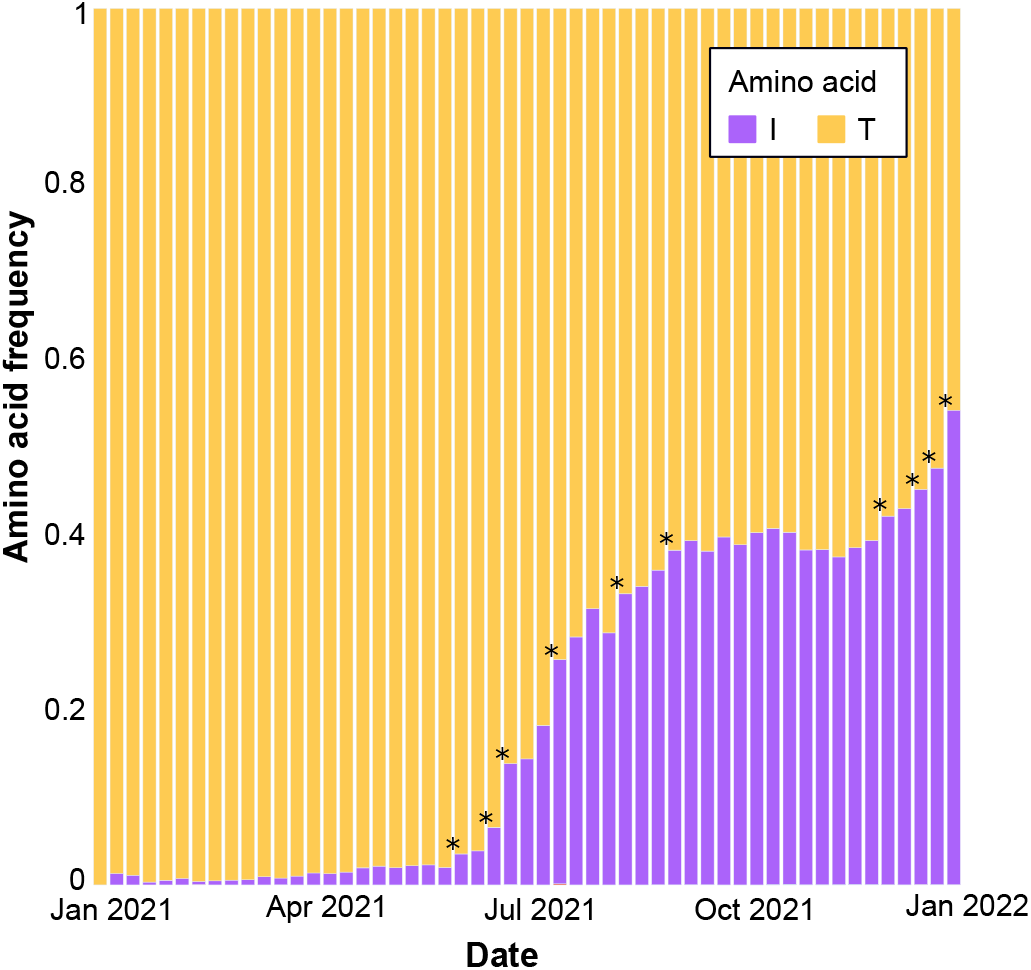
Frequency of amino acids at codon position 95 over calendar weeks (CW) for German sequences. The reference amino acid is threonine (T, yellow) and we can observe an increase in isoleucine (I, purple). Based on the underlying time series data downloaded from CovRadar, we calculated a significant increase of T95I in two time periods (CW21–34 and CW 47–52). Significant increases between consecutive CWs are marked by an asterisk and were calculated as described before [18].

### Front-end specifications

The CovRadar web service (covradar.net) uses the storage and compute resources of a dedicated Kubernetes (https://kubernetes.io/de/) cluster provided by the German Network for Bioinformatics Infrastructure (de. NBI). A MariaDB database, Redis caches, and web services are deployed in a redundant setup that allows for constant availability of the provided services. If necessary, the complete infrastructure setup can be easily scaled up on demand both in terms of storage volume and computing resources. All popular browsers are supported (see Table S2 for details). The database is hosted on local SSD storage to boost read performance. Database replication to ensure persistence is not needed as the database is working in a read-only-mode after the initial creation, and can be easily re-created in case of data loss. The whole setup is designed to provide a robust, scalable and performant infrastructure to run the CovRadar service.

## RESULTS

### CovRadar adds value via mutation-focused spike surveillance

During the pandemic, several tools were developed to analyze and visualize genomic and epidemiologic data to help track evolution and spread of the virus. Here, we collected web-based tools with a focus on SARS-CoV-2 data analysis and compared them to CovRadar’s features (Table S1). We want to briefly mention that several of the selected tools also provide additional functionalities, such as phylogenies and protein structures, that were not highlighted in CovRadar. We have developed CovRadar not to compete with existing platforms, but to add real value by focusing on individual mutations of the spike gene, which encodes an important target for vaccines. Thus, we focus here on a comparison of CovRadar’s features with the same features of the selected tools.

We collected 17 resources with a focus on their availability and SARS-CoV-2 relevance using Aviator [11], a tool for monitoring web services based on literature, OverCOVID (http://bis.zju.edu.cn/overcovid/resource.html), and our experience in daily work in SARS-CoV-2 surveillance. We excluded all non-molecular surveillance tools and non-working tools (although still online), resulting in this selection: GESS [12], Global evolution of SARS-CoV-2 [13], EMBL-EBI COVID-19 Data Portal [4], GISAID CoVSurver [6], CovidPhy [14], Coronavirus3D, NCBI, PathogenWatch, Nextstrain [8], CoV-GLUE [15], Outbreak, coVariants, CoV-Spectrum [10], COG-UK Mutation Explorer, COVID-19 variant dashboard, CoVerage, and Monitor of SARS-CoV-2 variants (Table S1). Most of these tools focus on information about lineages and their distribution over time, while only a few also provide distributions and spatial information for single mutations. In most cases, it is not possible to display information for different time periods on a map comparing multiple mutations. CoV-Spectrum is the only tool that provides a comparison with other variants and mutations, but it was not possible to compare different countries, calendar weeks or to select hosts. No tool offered alternative allele frequencies. In addition, no tool offered a built-in position converter, which is particularly useful when searching for mutations from the literature in an MSA. Other features are listed and compared in Table S1.

### Mutation frequency analyses on national level shows increasing frequencies of characteristic Omicron mutations within the last two weeks of December 2021

In this example, we investigate recent mutation dynamics within the spike gene with a focus on a specific country and time period. We use CovRadar’s filter options, allele frequency and mutation distribution plots, to show the rise of characteristic mutations in the spike gene in a short time frame and for the end of the year 2021 in Germany (Fig. 2 and 3). By taking delays in sampling, sequencing, and data submission into account, we restrict the data set to sequences from Germany of the last two calendar weeks in 2021 (CW51+52) and compare them to the consensus sequence from CW50. Thus, we only see the differences between consensus sequences specific for this time period and German sequences and not in comparison to Wuhan-Hu-1. Based on these filters, we can compare the AAF of 19,890 sequences (CW51+52) against the consensus based on 14,282 sequences (CW50) (Fig. 3).

Sites characteristic of Omicron (mutations with *>* 75% frequency in Omicron based on outbreak.info) show a higher AAF and are distributed among the different spike domains such as the Receptor Binding Domain (RBD) and the N-terminal domain (NTD) (see Fig. 2). The highest AAF at position 284 of the spike gene is within the NTD of the respective gene product (position 95 at the protein level). Taking a closer look on alternate alleles at this position (using the mutation distribution plot), we see that C has been replaced by T at position 284 which results in the T95I mutation profile on protein level (Fig. 3). The T95I change is characteristic for Omicron but has been previously described also for the Variant of Interest Mu and the Variant under Monitoring Iota and has been reported in vaccine breakthrough infections [16]. Furthermore, a pathogenicity analysis predicted T95I, among other mutations, to be deleterious in Omicron and sub-lineages [17]. We downloaded the complete time series data for C284T from the mutation distribution plot via the web app to perform an additional trend analysis. We compared the proportion of the alternative nucleotide T at position 284 between CWs using a Fisher exact test (adjusted p-value *<* 0.01, Table S3, see [18] for details on the method). We found that in two time periods (CW21-34 and CW 47-52), the proportion of samples with the alternative nucleotide T at position 284 increased significantly between consecutive weeks (Fig. 3). Based on these observations and analyses, we can expect T95I to become consensus, which is indeed the case in CW51 and subsequent weeks.

### Early emergence of spike change L452R in May 2021 in Germany

Using CovRadar, users can easily track spatio-temporal occurences of spike mutations, for example, linked to Variants of Concern (VOC). By default, we obtain information about characteristic VOC mutations from outbreak.info that can be selected by the user and then highlighted in a map. In this retrospective analysis, we investigate if characteristic Delta mutations were already evident in Germany in May 2021, a time where Alpha was still dominant. We select May 6 - 20, 2021 as the time frame and focus on the DESH data set with available postal codes. According to a weekly situation report by the Robert Koch Institute, Delta became dominant in Germany in CW25 (June, 2021) [19]. First, we selected N501Y which is strongly associated with VOC Alpha, Beta, and Gamma and E484K, known from Beta and Gamma. Both changes are only rarely observed in Delta (*<* 0.1%, outbreak.info). Second, we also selected L452R and T478K, which are both strongly represented in Delta, but only with low frequencies *<* 0.1% in Alpha, Beta, Gamma. The L452R change has been shown to increase infectivity *in vitro* and decrease neutralization by sera from COVID-19 patients and vaccinated individuals [20, 21] and has also been detected in other lineages such as A.27 which rose during the early months in 2021 in Germany [22]. The T478K change is commonly known from Delta and Omicron and has been predicted to affect the spike/ACE2 interaction [23]. Almost 100% of the sequences (21,128 / 21,156) that fall into the selected time frame (May 6 - 20, 2021) contain at least one of the four selected mutations (L452R, T478K, N501Y, E484K; Supplementary Fig. S3). We found that in May 2021, N501Y was dominant (Supplementary Fig. S3) but we can already observe a positive trend for other mutations such as L452R (Fig. 4) in some German regions.

**Figure 4.**
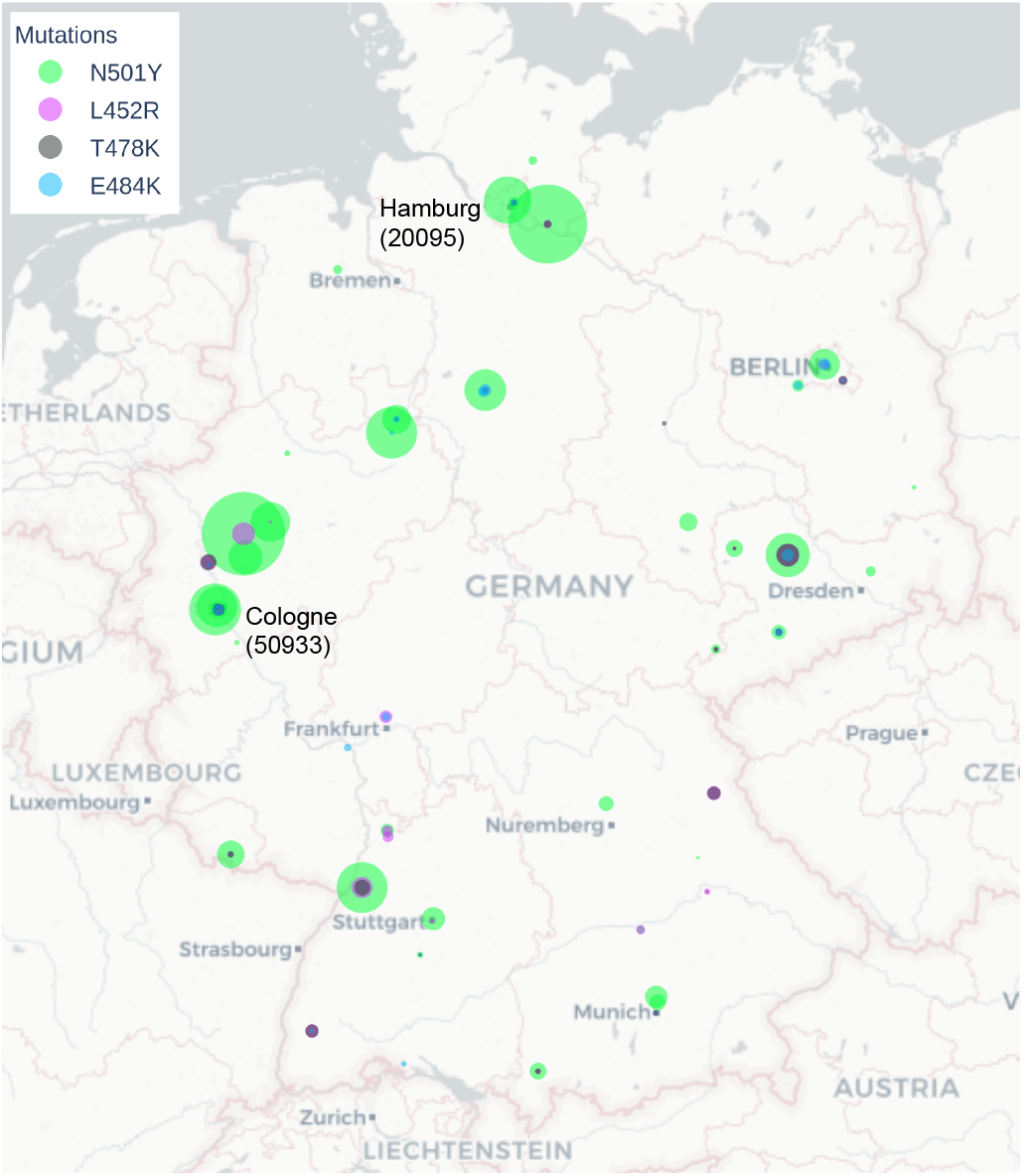
The geographic map shows the slope (circle size) of positive trends for the selected spike mutations L452R, T478K, E484K, and N501Y from May 6, 2021 to May 20, 2021. While N501Y is dominant throughout sampling locations (primary diagnostic labs) in Germany, we also observe a rise in other mutations such as L452R in certain areas.

To further investigate in which geographic location the first significant rise in L452R can be observed, we downloaded the selected time series data per location. We tested for significant differences in the proportion within the single postal code areas compared between calendar weeks (Fisher exact test, adjusted p-value *<* 0.01, Table S4, see [18] for details). We excluded data from February 2021 to rule out fluctuation effects due to increased sequencing capacity starting from January 25, 2021 in Germany. The first significant increase, excluding February, was sequenced in CW16 2021 (April 19–25 2021) for postal code 50933 (Cologne, North Rhine-Westphalia). In addition, we found that L452R significantly increased in CW31 with an 18 fold change in the postal code area 20095 (Hamburg). These examples also illustrate the importance of granular geographic data to track specific mutations and take action to contain new variants as quickly as possible. Please note that all sampling locations presented here are based on the primary diagnostic labs (performing the PCR test) and not the location of the patients due to data protection regulations.

## CONCLUSION

Molecular surveillance plays a crucial role in tracking the spread and evolution of SARS-CoV-2. Worldwide, millions of genome sequences have been generated and uploaded to databases such as GISAID and the COVID-19 Data Portal. While such data sources are treasure troves to answer various questions, the sheer amount of sequencing data directly leads to algorithmic and computational challenges, especially for non-bioinformaticians. Many web-based tools have been developed to provide fast access to already pre-processed data. However, these tools focus primarily on information about the SARS-CoV-2 lineage and Variants of Concern whose definition underlies a delay, while emerging Mutations of Concern are often neglected. Thus, we present CovRadar, a web service to enhance molecular surveillance of the spike gene with a focus on the mutation level and customizable spatial and time analyses. We believe that CovRadar will aid researchers around the world to get better and faster access to SARS-CoV-2 mutation profiles and metadata to support the ongoing fight against the COVID-19 pandemic on the molecular surveillance level.

## AVAILABILITY

CovRadar is freely accessible at https://covradar.net, its open-source code is available at https://gitlab.com/dacs-hpi/covradar.

## SUPPLEMENTARY DATA

Supplementary Data are available at NAR online.

## ACKNOWLEDGEMENTS

Many thanks to Elizabeth Yuu for proofreading the manuscript. We are very grateful to the GISAID Initiative, the COVID-19 Data Portal hosted by EMBL-EBI, the German Electronic Sequence Data Hub (DESH, https://github.com/robert-koch-institut/SARS-CoV-2-Sequenzdaten_aus_Deutschland) and all data contributors, i.e. the authors from the originating laboratories responsible for obtaining the specimens and the submitting laboratories where genetic sequence data were generated and shared and on which this research is based.

