## Supplementary Material for "CovRadar: Continuously tracking and filtering SARS-CoV-2 mutations for molecular surveillance"

#### This PDF file includes:

Supplementary Methods
Figs. S1 to S3
Tables S1 to S4
SI References

### Supplementary Methods

#### 1. Analytical pipeline

The analytical back-end of CovRadar, including input data pre-processing and merging of different data sources, is implemented in a Snakemake (1) pipeline utilizing further custom Python scripts to enable reproducible, expandable, and resumable analyses (gitlab.com/dacs-hpi/covradar, "backend" directory).

Below, we explain in detail the algorithms and third-party tools used in the analytical workflow to generate the results displayed on CovRadar's web application. An overview of the analytical pipeline is shown in Fig. S1, while further details of the algorithms are described in the following subsections.

**A. Data Preprocessing.** CovRadar needs as input SARS-CoV-2 genome sequences in compressed multi-FASTA format and corresponding metadata in TSV format. The metadata can comprise information such as sampling date, host, country, postal code, and lineage assignments as provided by GISAID (2–4) or the COVID-19 Data Portal hosted by EMBL-EBI (5). CovRadar can integrate different data sources including third-party data sets which can be beneficial if those different sources provide complementary data. For the back-end analysis pipeline, minimal metadata comprises country, sampling date, and host. To obtain the full functionality of the web application, additional information on lineage assignment and postal codes are required. If no postal codes are available, country information will be used. The online instance of CovRadar comprises the COVID-19 Data Portal data set automatically downloaded via API access. Alternatively, data provided by GISAID can be automatically downloaded and used as data source if API access is available. Every laboratory in Germany that sequences SARS-CoV-2 is required by the Coronavirus Surveillance Regulation (CorSurV) of the Federal Ministry of Health to transmit the resulting genome sequences and associated metadata to the Robert Koch Institute. This transmission is done via the German Electronic Sequence Data Hub (DESH). Data curation and quality management of the (meta-)data was done by the Robert Koch Institute before sharing the data publicly (github.com/robert-koch-institut/SARS-CoV-2-Sequenzdaten_aus_Deutschland) and Zenodo (doi.org/10.5281/zenodo.5139363). Since the DESH data set provides more detailed metadata such as postal codes of the respective primary diagnostics laboratory and regularly updated virus lineage assignments, it is considered as well. To prevent duplicates in the data collection, we remove all German sequences from EMBL-EBI and merge the data sets with the DESH data for further analysis.

**B. Files extraction.** CovRadar accepts as input compressed FASTA files containing the genomes and TSV files with their respective metadata. The file extraction is automated by a shell script capable of recognizing and extracting multiple formats, namely: 7z (.7z), bzip2 (.bz2), gzip (.gz), RAR (.rar), TAR (.tar), TBZ2 (.tar.bz2 or .tbz2), TGZ (.tar.gz or .tgz), Z (.Z) and zip (.zip).

**C. Merging of data sets.** CovRadar's pipeline accepts multiple different sources as input. For example, it is possible to simultaneously use EMBL-EBI, GISAID, and other third-party data sets. This can be beneficial if those different sources have complementary data. The data sets merging is performed by a Python script that accepts as input only DNA sequences with IUPAC nucleotide codes (6). Any sequences with unrecognized characters are removed to avoid errors in further steps of the workflow. File handling is leveraged by Biopython (7) and pandas (8) for the FASTA and TSV files, respectively.

**D. Spikes extraction.** In order to extract the spikes, first the genomes are aligned to the spike of Wuhan-Hu-1 (NC_045512.2), the first case from Wuhan. This alignment is performed by pblat (9), using default parameters. The software pblat is a scalable implementation of the popular aligner blat, which is especially useful in cases where other aligners may have trouble, e.g., when sequences are too long or the alignments have large gaps (9). Afterwards, an in-house Python script extracts the spike sequences from the genomes according to the coordinates detected by pblat, keeping only alignments with an N content smaller than or equal to 5 % and a length deviation also smaller than or equal to 5 % when compared to the spike from Wuhan-Hu-1. Filtering the spikes this way allows for an easy exclusion of sequences with low quality or with assembly errors. This script also employs pandas (8) and Biopython (7) to manipulate the TSV and FASTA files. The extraction Python script can be also used stand-alone and for other genes.

**E. Multiple sequence alignment.** The codon-aware multiple sequence alignment (MSA) of the extracted spike sequences is performed by VIRULIGN (10) using default parameters. The spike from Wuhan-Hu-1 is used as a reference and the results are exported as a global alignment composed of nucleotides.

**F. Computation of consensus sequences.** Before the metadata-specific consensus sequences are computed, first the spike from Wuhan-Hu-1 is extracted from the MSA produced by VIRULIGN, in order to be kept as reference. Afterwards, the MSA's sequences are divided by country and calendar weeks to allow for easier interpretation of the results.

The calendar weeks are computed using the ISO standard provided by the Python library isoweek, while sequences with incomplete or invalid dates are removed to keep the integrity of the results. Finally, the consensus sequences are computed from those divided subsets. The consensus algorithm counts how many A, T, C, G or gaps (-) appear per position, keeping whichever has the most occurrences or alphabetical order in case of ties. If none of those options arise, then an N is inserted in that position, thus, accounting for ambiguous base pairs.

If the consensus nucleotide at a given position differs from the reference, we translate the resulting codon and display the corresponding amino acid. As VIRULIGN is codon-aware, gaps are translated into missing amino acids (-). Each substitution is treated separately, so that its effect on the amino acid sequence can be examined independently from other mutations. However, we have not observed multiple substitutions occurring within the same codon of the consensus sequence yet.

**G. Numbering.** To facilitate comparative results, we have introduced a numbering system based on Wuhan-Hu-1. With this numbering, each MSA position can be converted to the corresponding position of the first case. The script takes the aligned first case. Since the unaligned first case has no gaps, an alignment will at most result in a longer sequence. Each gap corresponds to one insertion. For deletions, the MSA will contain the gaps, but the length of the first case will not change. To get the corresponding positions of Wuhan-Hu-1, the script increments the position at every base that is not a gap. If there is a gap, the last position is taken instead with a suffix ".X", where X stands for an integer. The result is stored in a TSV table.

**H. Variant counting.** For obtaining variants (nucleotide substitutions, INDELs), the MSA and the aligned Wuhan-Hu-1 sequence are required as input. The first case will be the reference column (REF) in the VCF file. The script goes through every position in the MSA and adds differences as alternative alleles (ALT). The position refers to the MSA and is stored in the column POS. Only positions with variations are stored in the VCF file. Due to performance reasons, first, each row of the VCF file is saved in a temporarily directory. After each row is completed, the rows are then merged into one VCF file.

**I. Mutation profile assignment.** To display the mutations in the map of the web application, we create with custom Python scripts links between all sequences and the mutations they carry. Thereby the mutations are provided as a TSV file as input data. They can be of any mutation (substitutions, INDELs) or combinations of them. In the repository, we provide a file that contains all characteristic spike mutations that occur in over >75% of current VOC, based on outbreak.info.

**J. Basic statistics.** Basic statistics are computed with BCFtools (11, 12) and RAxML-NG (13). For BCFtools the stats parameter is used, while for RAxML-NG we use the parameter --check. As a best-fitting nucleotide substitution model and evolutionary model, a generalized time-reversible plus gamma distribution (+G) was chosen by ModelTest-NG (14). We specifically optimized the base frequencies by maximum-likelihood (+FO).

**K. Plots creation.** The pipeline also creates a PDF report for a first overview. The PDF reports contain two charts, one representing the nucleotide substitutions and the other showing the allele frequencies. The substitutions are computed by BCFtools, which stores its results in text format. Those results are parsed with the Python library *re* using regular expressions, then they are plotted with the library *matplotlib*. The PDF reports use a modified allele frequency algorithm of the web application to create the allele frequency plot. Please see Section 2. E for a detailed explanation and the differences.

**L. Preliminary analyses.** CovRadar offers stand-alone scripts to analyse the output of the Snakemake pipeline directly if setting up a database and hosting a web service are not possible or required.

**M. Creating the SQL dump.** All results provided by the analytical pipeline are imported into a MariaDB database for the web service using a blue-green deployment to ensure continuous access. Thereby, the data from the COVID-19 data portal (covid19dataportal.org) (updated bi-weekly) and the reduced data set with the sequences of the last four weeks (updated every other day) each receive their own database. The SQL dumps are then uploaded to the Kubernetes cluster run and hosted by the German Network for Bioinformatics Infrastructure, de.NBI.

### 2. Web application

Below is additional information about the web application that is not included in the main script.

**A. Browser compatibility.** For browser compatibility, please refer to Table S2.

**B. Layout.** A landing page guides the user to the web applications: the full data set (updated bi-weekly) and the most recent data set of the last four weeks (updated daily). Further links to the publication, source code, supporters, and a help page are provided. The web application itself is split into three main panels: right menu, left menu, and a middle panel showing the actual content. The right menu is used for navigation through the content of the app and provides a switch between English and German text as well as a link to the help page. We implemented a position converter to easily translate between nucleotide and amino acid positions as well as codons in the MSA and in regard to the reference spike gene (further details below). The left menu provides different sequence filters specifically for the content shown in the middle panel, such as host, time range, and country. The middle panel shows the interactive plots and tables with direct access to downloadable figures and the raw data in TSV format. Each plot is equipped with an information button explaining the content and provides content-specific interactive tooltips via mouse-over and zoom functionalities.

**C. Help Page.** The help page (covradar.net/help) can be accessed from the landing page and the right menu of the app. It also provides detailed and illustrated descriptions of every functionality, the different plots, the tables as well as information on compatible browsers and a FAQ.

**D. General data characteristics and alignment statistics.** A data distribution plot provides an overview of the available SARS-CoV-2 genome sequences per calendar week and country and allows to identify over-/ underrepresented countries in the data set. The "global" option in the web application always refers to all sequences included in the data set. A statistics panel provides additional information about the data source and pipeline as well as alignment metrics. We show when the analytical pipeline started the last time and based on which data sources. A simplified overview of the analytical back-end helps put the different results into perspective. The analytical pipeline also calculates alignment metrics using RAxML-NG and BCFtools to help investigate the quality of the MSA which underlies all further reports. We provide warnings about duplicated sequences with the exact same sequence and provide numbers about the percentage of gaps in the MSA and the number of detected variants and multiallelic sites.

**E. Alternative allele frequency plot.** The allele frequency plot shows the frequency of alternative alleles in relation to Wuhan-Hu-1 or a selected consensus sequence. (example Fig. 2 in main script) For the frequencies of the alternative alleles of a requested subsample, these sequences are filtered out of the VCF file using country, start date, end date, origin and host. The reference in the VCF file is Wuhan-Hu-1. Thus, it must be taken into account if the user has selected a consensus sequence against which the frequencies should be calculated. For this purpose the positions that show differences between the consensus sequence and Wuhan-Hu-1 are determined and it is checked if they show alternative alleles in the VCF file. If not, all sequences for this position have the base of the first case. This means that in relation to the consensus base at this position the frequency of the alternative alleles is 1. For all other positions, where the first case base and the consensus differ, the alternative alleles are identified again. For some cases no clear statement can be drawn if an alternative allele is present because, in contrast to Wuhan-Hu-1 and the consensus sequences, degenerated bases can occur in the sequences. For example, this happens if there is only information that the sequence base is purine and the reference base is adenine or guanine. In this case, only pyrimidines are counted as alternative alleles and otherwise excluded. Finally, for each position the number of non-excluded alternative bases is divided by the number of non-excluded sequences at that position and the result is returned as a list of frequencies. Since only non-degenerated bases as well as gaps were used in the consensus sequence, it may occur that there is no coverage for a position. In this case, the consensus sequence shows the base N. If the consensus base is N, no frequency is determined.

**F. Consensus sequence table.** The consensus sequence table (Fig. S2) reports changes in the consensus compared to the Wuhan-Hu-1 reference sequence (NC_045512.2) for each calendar week and chosen country. Each row of the table shows the mutational differences between the consensus sequence of that calendar week in comparison to the reference. The table header shows the position with respect to the MSA as well as with respect to the reference. The first row contains the corresponding reference bases with associated amino acid translations. If the consensus base matches the reference base, it is marked with a dot. Deletions are marked by a dash, "-". Next to the calendar week label, the sequence coverage is given. For each position, the number of sequences supporting that base and the translated amino acid (via the position converter script) are provided. In case of insertions, the reference shows a gap "-" and the inserted nucleotides are given for the consensus.

***F.1. Comment on amino acid translations.*** It is possible that two nucleotide substitutions, each synonymous by itself, can alter the amino acid when they coincide. For example: CTA codon (Leu) still codes for Leu if changed to CTC and TTA, but both combined result in a TTC codon translating into Phe. Similarly, it is possible that two nucleotide substitutions are non-synonymous but lead to a different amino acid change when they coincide. For example: TGG (Trp) may change to TGC (Cys) or TTG (Leu), but if combined the two substitutions result in the TTC codon coding for Phe. In principle, it is possible to translate all the sequences for a given calendar week individually to built a consensus protein sequence. However, it is possible that the consensus amino acid at a given position can be different from the one displayed based on a single consensus nucleotide substitution. Please see Section 1.F for further details on the algorithm.

**G. Caching.** CovRadar's analytical pipeline can handle a large number of sequences (over 4 million sequences for EMBL-EBI in March 2022), which then leads to web application response time issues due to complex database queries and the sheer size of the underlying database. To solve this issue, we cache slow calculations like the initial load of the web application and additional settings that are used often in Dash callbacks. In particular, we store the final HTML results of slow SQL queries that are otherwise too big to cache in a very fast in-memory database (github.com/redis/redis). Thus, we are able to reduce initial loading times from one minute to less than ten seconds. When the database or web application is updated, the cache is no longer valid which would result in sub-optimal user experiences when first accessing the page. To prevent this, we automatically visit the web page during the deployment and activate commonly used features via Selenium (selenium.dev) to cache them.

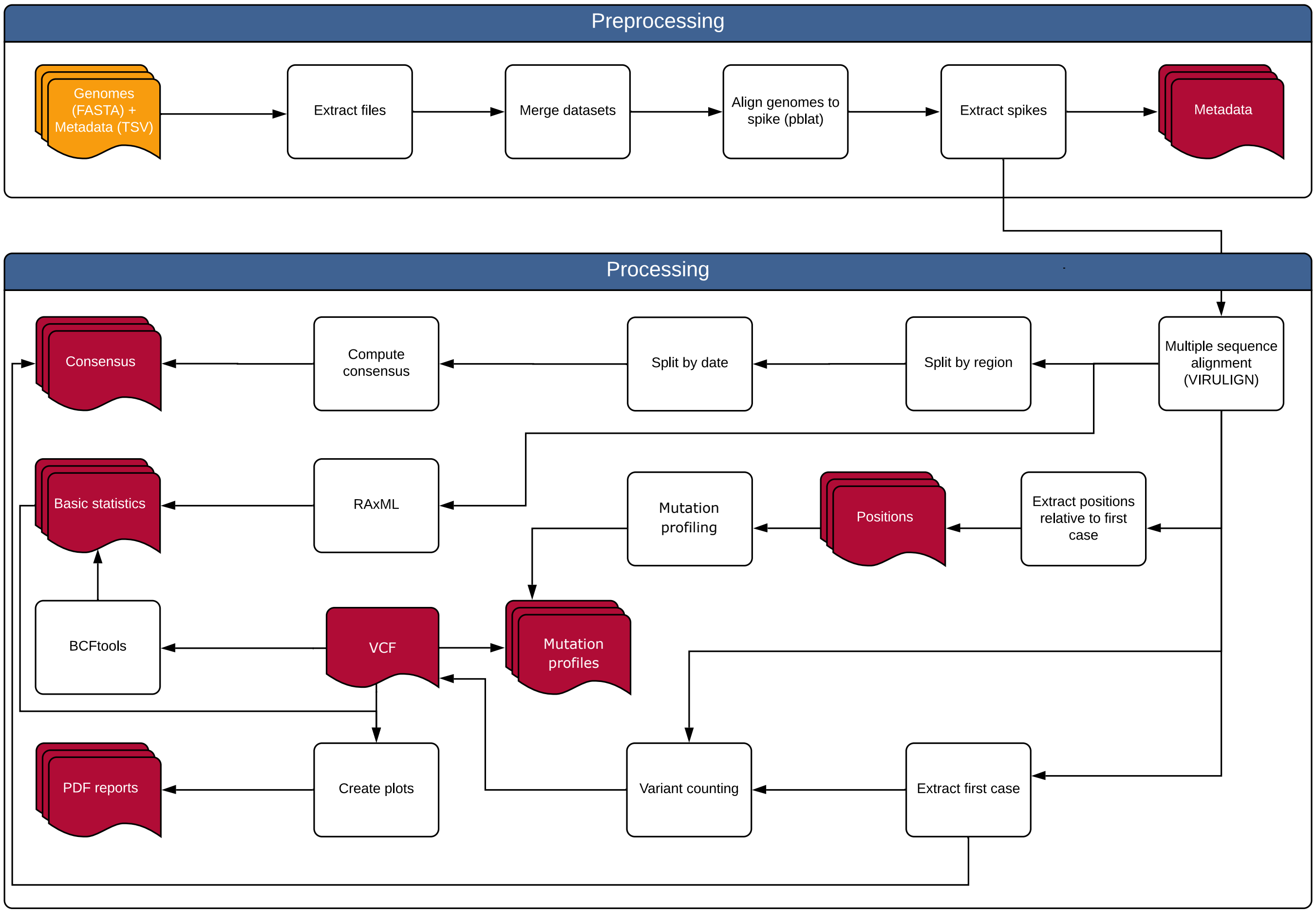


**Fig. S1.** Analysis workflow used to generate the results displayed on CovRadar's website. Orange blocks depict input files, while white blocks are processing steps of the pipeline and red blocks represent output that is displayed on the website. The pipeline accepts as input compressed FASTA files containing the sequences and TSV files with their metadata. One or more different data sources can be used simultaneously, e.g., EMBL-EBI COVID-19 Data Portal and DESH (national data from the German Electronic Sequence Data Hub). First the pipeline extracts the files, then merges the data sets if more than one is used. Afterwards, pblat is used to align the input genomes against the spike sequence from Wuhan-Hu-1. Then the spikes are extracted with the coordinates reported by pblat. Next, VIRULIGN is used to perform a codon aware multiple sequence alignment (MSA) of the extracted spikes. Before computing the consensus sequences from the MSA, the sequences are first separated by country and calendar week. Additionally, the Wuhan-Hu-1 is extracted from the MSA and added to the consensus sequences to be used as reference. The coordinates relative to Wuhan-Hu-1 are extracted from the MSA with an in-house script. Finally, the variant counting is performed by an in-house script and basic statistics are computed with RAxML and BCFtools. In addition, a mutation profile is generated based on the VCF for each sequence.

| msa | | 1203 | | | 1204 | | | 5549 | | | 5858 | | | 6007 | | | 6449 | | | 7331 | | | 7694 | | | 7838 | | | 8896 | | | 9001 | | |
|---|---|---|---|---|---|---|---|---|---|---|---|---|---|---|---|---|---|---|---|---|---|---|---|---|---|---|---|---|---|---|---|---|---|---|
| ref | | 471 | | | 472 | | | 1355 | | | 1433 | | | 1501 | | | 1709 | | | 1841 | | | 2042 | | | 2147 | | | 2848 | | | 2944 | | |
| Wuhan-Hu-01 | 1 | C | F157F | 1 | A | R158R | 1 | T | L452L | 1 | C | T478T | 1 | A | N501N | 1 | C | A570A | 1 | A | D614D | 1 | C | P681P | 1 | C | T716T | 1 | G | D950D | 1 | T | S982S | 1 |
| 2021W26 | 1452 | - | - | 1005 | G | R158G | 987 | G | L452R | 1019 | A | T478K | 999 | . | . | 1024 | . | . | 1108 | G | D614G | 1450 | G | P681R | 999 | . | . | 1110 | A | D950N | 721 | . | . | 1110 |
| 2021W25 | 1379 | - | - | 687 | G | R158G | 684 | G | L452R | 740 | A | T478K | 704 | . | . | 746 | . | . | 837 | G | D614G | 1378 | G | P681R | 702 | . | . | 827 | . | . | 689 | . | . | 830 |
| 2021W24 | 1922 | . | . | 1183 | . | . | 1185 | . | . | 1134 | . | . | 1171 | T | N501Y | 1077 | A | A570D | 984 | G | D614G | 1920 | A | P681H | 1024 | T | T716I | 984 | . | . | 1187 | G | S982A | 979 |
| 2021W23 | 3184 | . | . | 2727 | . | . | 2728 | . | . | 2643 | . | . | 2725 | T | N501Y | 2542 | A | A570D | 2418 | G | D614G | 3174 | A | P681H | 2471 | T | T716I | 2435 | . | . | 2721 | G | S982A | 2429 |
| 2021W22 | 4650 | . | . | 4300 | . | . | 4304 | . | . | 4207 | . | . | 4296 | T | N501Y | 4069 | A | A570D | 3953 | G | D614G | 4618 | A | P681H | 3977 | T | T716I | 3950 | . | . | 4309 | G | S982A | 3946 |
| 2021W21 | 5845 | . | . | 5598 | . | . | 5591 | . | . | 5517 | . | . | 5587 | T | N501Y | 5383 | A | A570D | 5187 | G | D614G | 5794 | A | P681H | 5237 | T | T716I | 5197 | . | . | 5604 | G | S982A | 5175 |
| 2021W20 | 8042 | . | . | 7849 | . | . | 7848 | . | . | 7741 | . | . | 7851 | T | N501Y | 7571 | A | A570D | 7381 | G | D614G | 7991 | A | P681H | 7400 | T | T716I | 7382 | . | . | 7867 | G | S982A | 7370 |

**Fig. S2.** CovRadar's consensus table showing differences of global consensus sequences of single calendar weeks compared to the Wuhan-Hu-1 reference sequence. The table contains position, consensus base and coverage of changes in the consensus base regarding to Wuhan-Hu-1. The table header shows the position with respect to the multiple sequence alignment (msa) or with respect to Wuhan-Hu-1 (ref). The row starting with "first-case" contains the corresponding bases of the first case (Wuhan-Hu-1). Below are the consensus bases per calendar week. If they match the first case, they are marked with a dot. Gaps are indicated with "-", insertions with "ins". Next to the calendar week is the sequence coverage for that week. Next to the bases is the number of sequences that have this base at this position. Along with the nucleotide mutation, the associated amino acid mutation is listed. Data were obtained from the EMBL-EBI database.

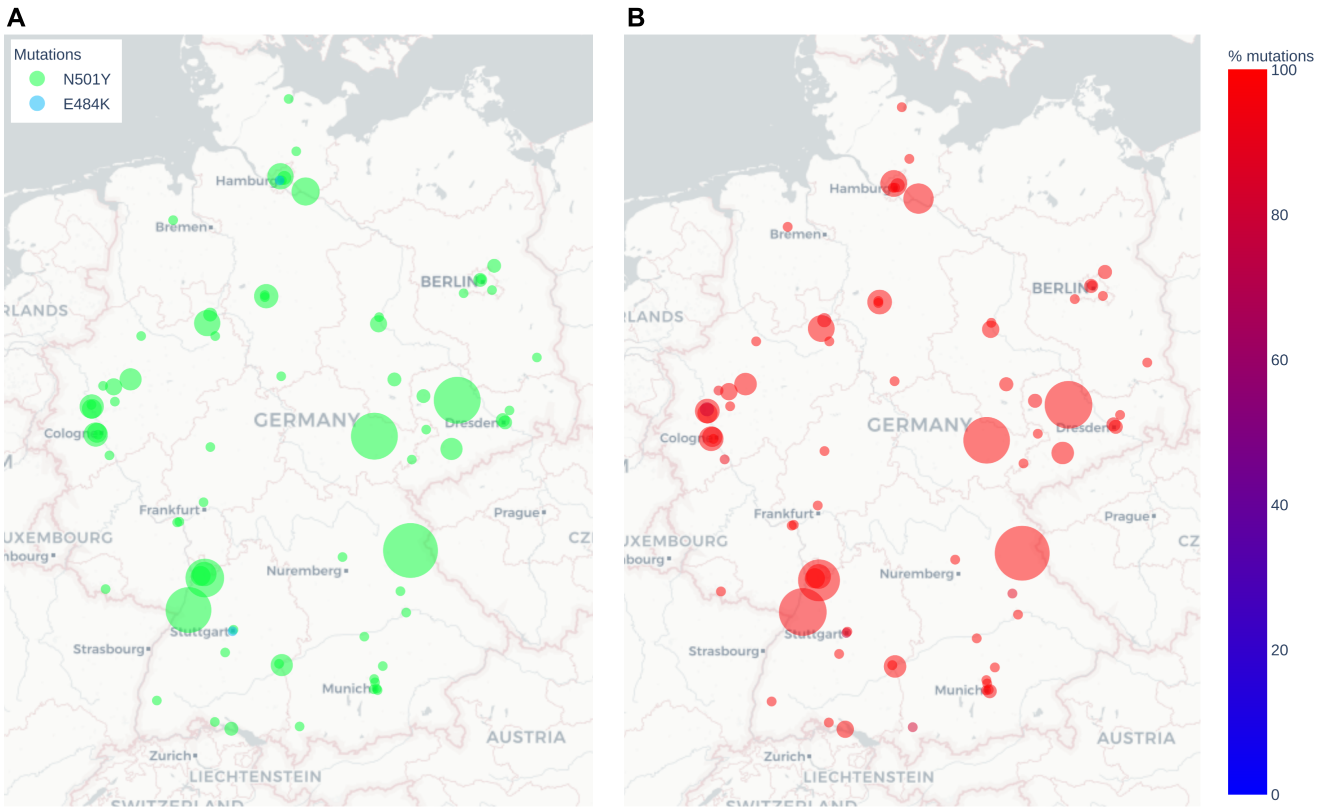


**Fig. S3.** **A**: The geographic map shows the most frequent mutation per sampling location (primary diagnostic labs) during May 6 – May 20, 2021 with frequencies represented by circle size. Spike change N501Y commonly known from Alpha, Beta, and Gamma is dominant, **B**: The proportion of sequences containing one of the following selected spike changes: L452R, T478K, E484K, and N501Y with frequencies represented by circle size. Out of 21,156 sequences in the selected time period 21,128 have at least one of the four mutations, thus, our mutation selection allows representation for sequences during that time period in Germany.

**Table S1. Table comparing CovRadar's features with other tools. Note that we focused on features that are provided by CovRadar. Many tools include other helpful features such as phylogeny or protein structure, which is not the focus of CovRadar. "x": present, "-": absent, "1": present but not for every nucleotide site in the RBD, "2": present but not for every country, "3": information present in table but not in plot. nt – nucleotide level; aa – amino acid level; CW – calender week. URLs last accessed March 7th, 2022.**

| Name$^{URL}$ (citation if available) | Map with aa mutation occurrences over time | Alternative frequencies based on consensus sequences per country and CW | Non-Wuhan consensus sequences per CW per country | Mutation distribution plot for every country – nt | Mutation distribution plot for every country – aa | Position converter | Based on mutations | Provided data download in PNG and CSV format | Global data |
|---|---|---|---|---|---|---|---|---|---|
| **CovRadar**[1] | x | x | x | x | x | x | x | x | x |
| GESS[2] (15) | - | - | - | 1,2,3 | - | - | x | x | x |
| Global evolution of SARS-CoV-2[3] (16) | - | - | - | - | - | - | - | - | x |
| EMBL-EBI COVID-19 Data Portal[4] (17) | - | - | - | - | - | - | - | x | x |
| GISAID CoVSurver[5] (4) | - | - | - | - | 1 | - | x | x | x |
| covidPhy[6] (18) | - | - | - | - | - | - | - | - | x |
| coronavirus3d[7] | - | - | - | - | 3 | - | x | x | x |
| NCBI[8] | - | - | - | - | - | - | x | - | x |
| PathogenWatch[9] | - | - | - | - | - | - | - | x | x |
| Nextstrain[10] (19) | x | - | - | x | x | - | x | x | x |
| CoV-GLUE[11] (20) | - | - | - | - | - | - | x | - | x |
| Outbreak[12] | - | - | - | - | x | - | x | - | x |
| coVariants[13] | - | - | - | - | 1 | - | x | - | x |
| CoV-Spectrum[14] (21) | x | - | - | - | x | - | x | x | x |
| COG UK mutation explorer[15] | - | - | - | - | x | - | x | x | - |
| COVID-19 variant dashboard[16] | - | - | - | - | - | - | - | - | - |
| CoVerage[17] | - | - | - | - | 1,2 | - | x | - | x |
| Monitor of SARS-CoV-2 variants[18] | - | - | - | - | - | - | - | - | x |

**URLs**
[1] https://covradar.net
[2] https://wan-bioinfo.shinyapps.io/GESS
[3] http://www.covid19evolution.net
[4] https://www.covid19dataportal.org
[5] http://www.gisaid.org/epiflu-applications/covsurver-mutations-app
[6] http://covidphy.eu
[7] https://coronavirus3d.org
[8] https://www.ncbi.nlm.nih.gov/labs/virus/vssi/#/scov2_snp
[9] https://pathogen.watch/collection/1y3cdsjq55hf-public-genomes
[10] https://nextstrain.org/ncov/gisaid/global
[11] http://cov-glue.cvr.gla.ac.uk
[12] outbreak.info
[13] https://covariants.org
[14] https://cov-spectrum.org
[15] https://sars2.cvr.gla.ac.uk/cog-uk
[16] https://gis.ecdc.europa.eu/portal/apps/opsdashboard/index.html#/25b6e879c076412aaa9ae7adb78d3241
[17] https://sarscoverage.org
[18] https://monitor.mi2.ai

**Table S2. Tested operating systems and browsers with version number. Note, that these are the browser versions we specifically used for testing. Older versions will likely also work. Mobile browsers and Internet Explorer are not supported.**

| Operating System | Firefox | Chrome | Edge | Safari |
|---|---|---|---|---|
| Windows 10 | 90 | 90 | 90 | n/a |
| macOS Monterey | 90 | 96 | 90 | 15 |
| Ubuntu 20.04 | 93 | 96 | n/a | n/a |

**Table S3. Table showing significant increase between two calendar weeks for position 284 (codon 95). C284T (amino acid S:T95I) had significantly increased for the first time by 1.8 fold in CW21 2021. "nt": nucleotide, "Y": C or T, "cw": calendar week, "conf.low": lower border of confidence interval, "conf.high": higher border of confidence interval, "p.adj": adjusted p value**

| nt | YEAR | CW | fold change | p.value | conf.low | conf.high | p.adj |
|---|---|---|---|---|---|---|---|
| C | 2021 | 21 | 0,57 | 5,05E-08 | 0,47 | 0,70 | 1,03E-06 |
| C | 2021 | 23 | 0,58 | 7,28E-08 | 0,47 | 0,71 | 1,39E-06 |
| C | 2021 | 24 | 0,44 | 2,52E-18 | 0,36 | 0,53 | 1,45E-16 |
| C | 2021 | 26 | 0,69 | 1,47E-04 | 0,56 | 0,84 | 1,40E-03 |
| C | 2021 | 27 | 0,69 | 1,36E-05 | 0,58 | 0,82 | 1,69E-04 |
| C | 2021 | 31 | 0,85 | 4,58E-04 | 0,77 | 0,93 | 4,10E-03 |
| C | 2021 | 34 | 0,90 | 5,19E-04 | 0,85 | 0,96 | 4,51E-03 |
| C | 2021 | 47 | 0,90 | 5,45E-05 | 0,86 | 0,95 | 5,59E-04 |
| C | 2021 | 50 | 0,93 | 1,25E-03 | 0,88 | 0,97 | 9,43E-03 |
| C | 2021 | 51 | 0,75 | 1,59E-29 | 0,72 | 0,79 | 1,52E-27 |
| C | 2021 | 52 | 0,60 | 3,36E-65 | 0,56 | 0,63 | 9,65E-63 |
| T | 2021 | 21 | 1,80 | 3,95E-08 | 1,45 | 2,23 | 8,73E-07 |
| T | 2021 | 23 | 1,73 | 1,85E-07 | 1,40 | 2,13 | 3,12E-06 |
| T | 2021 | 24 | 2,29 | 1,25E-17 | 1,89 | 2,79 | 5,96E-16 |
| T | 2021 | 27 | 1,54 | 1,18E-06 | 1,29 | 1,84 | 1,88E-05 |
| T | 2021 | 31 | 1,23 | 1,96E-05 | 1,12 | 1,36 | 2,35E-04 |
| T | 2021 | 34 | 1,11 | 1,02E-03 | 1,04 | 1,17 | 7,95E-03 |
| T | 2021 | 47 | 1,12 | 3,24E-06 | 1,07 | 1,18 | 4,66E-05 |
| T | 2021 | 49 | 1,09 | 4,46E-04 | 1,04 | 1,14 | 4,10E-03 |
| T | 2021 | 50 | 1,10 | 5,38E-05 | 1,05 | 1,16 | 5,59E-04 |
| T | 2021 | 51 | 1,30 | 3,52E-26 | 1,24 | 1,37 | 2,52E-24 |
| T | 2021 | 52 | 1,50 | 2,27E-43 | 1,42 | 1,60 | 3,25E-41 |
| Y | 2021 | 38 | 4,17 | 1,83E-07 | 2,27 | 8,15 | 3,12E-06 |
| Y | 2021 | 39 | 0,31 | 2,10E-05 | 0,16 | 0,56 | 2,41E-04 |
| Y | 2021 | 40 | 4,36 | 1,36E-08 | 2,45 | 8,26 | 3,26E-07 |
| Y | 2021 | 47 | 1,98 | 7,10E-04 | 1,31 | 3,05 | 5,66E-03 |

**Table S3. Table showing significant differences in the proportion within the single German postal codes (ZIP) compared between calendar weeks (CW) for S:L452R. The first significant increase, excluding February, was sequenced in CW16 2021 for ZIP 50933 (Cologne, North Rhine-Westphalia). zip – German postal code, CW – calendar week, conf.low – lower border of confidence interval, conf.high – higher border of confidence interval, p.adj – adjusted p value, Inf – fold change cannot be estimated.**

| zip | YEAR | CW | fold change | p.value | conf.low | conf.high | p.adj |
|---|---|---|---|---|---|---|---|
| 76131 | 2021 | 6 | 11,05 | 1,77E-198 | 9,33 | 13,11 | 5,32E-195 |
| 44879 | 2021 | 8 | 12,81 | 4,25E-37 | 8,02 | 21,25 | 5,31E-35 |
| 50933 | 2021 | 16 | Inf | 1,59E-06 | 4,86 | Inf | 2,46E-05 |
| 07745 | 2021 | 17 | 8,72 | 2,60E-05 | 2,58 | 46,04 | 3,28E-04 |
| 69214 | 2021 | 22 | Inf | 1,13E-03 | 2,23 | Inf | 9,14E-03 |
| 69126 | 2021 | 23 | 4,52 | 4,18E-05 | 2,02 | 11,49 | 4,89E-04 |
| 32105 | 2021 | 25 | 4,29 | 2,08E-04 | 1,82 | 11,76 | 2,06E-03 |
| 20251 | 2021 | 27 | 7,98 | 3,53E-08 | 3,16 | 25,89 | 7,51E-07 |
| 13353 | 2021 | 28 | 1,98 | 8,97E-04 | 1,30 | 3,08 | 7,40E-03 |
| 78467 | 2021 | 29 | 3,74 | 4,56E-10 | 2,33 | 6,28 | 1,20E-08 |
| 04779 | 2021 | 29 | 3,25 | 6,79E-05 | 1,71 | 6,71 | 7,56E-04 |
| 76131 | 2021 | 29 | 1,92 | 1,66E-10 | 1,56 | 2,39 | 4,57E-09 |
| 40477 | 2021 | 29 | 1,59 | 1,09E-04 | 1,25 | 2,03 | 1,15E-03 |
| 70174 | 2021 | 30 | 3,23 | 1,16E-04 | 1,67 | 6,75 | 1,22E-03 |
| 50858 | 2021 | 30 | 2,03 | 9,70E-05 | 1,39 | 2,99 | 1,04E-03 |
| 20095 | 2021 | 31 | 17,68 | 1,61E-18 | 6,68 | 66,23 | 8,61E-17 |
| 22081 | 2021 | 31 | 7,04 | 1,02E-08 | 3,05 | 19,97 | 2,29E-07 |
| 50858 | 2021 | 31 | 3,76 | 1,21E-06 | 2,04 | 7,46 | 1,93E-05 |
| 44879 | 2021 | 31 | 3,53 | 3,63E-04 | 1,63 | 8,74 | 3,40E-03 |
| 04779 | 2021 | 31 | 1,82 | 4,26E-19 | 1,59 | 2,08 | 2,41E-17 |
| 50858 | 2021 | 31 | 1,57 | 7,50E-10 | 1,35 | 1,82 | 1,92E-08 |
| 66123 | 2021 | 32 | 18,13 | 1,02E-09 | 4,72 | 154,43 | 2,56E-08 |
| 50858 | 2021 | 32 | 12,93 | 5,30E-04 | 2,07 | 535,06 | 4,66E-03 |
| 50935 | 2021 | 32 | 11,84 | 7,79E-04 | 1,96 | 483,29 | 6,54E-03 |
| 40477 | 2021 | 32 | 2,88 | 2,41E-04 | 1,57 | 5,63 | 2,34E-03 |
| 56068 | 2021 | 32 | 2,74 | 1,12E-13 | 2,05 | 3,69 | 4,20E-12 |
| 78467 | 2021 | 32 | 2,53 | 4,76E-05 | 1,57 | 4,21 | 5,50E-04 |
| 21502 | 2021 | 32 | 1,27 | 6,41E-05 | 1,13 | 1,43 | 7,27E-04 |
| 20251 | 2021 | 33 | 13,73 | 2,77E-04 | 2,25 | 562,72 | 2,64E-03 |
| 92637 | 2021 | 33 | 5,14 | 1,14E-05 | 2,19 | 14,73 | 1,53E-04 |
| 21502 | 2021 | 33 | 3,46 | 2,96E-04 | 1,64 | 8,16 | 2,81E-03 |
| 48143 | 2021 | 33 | 2,91 | 6,36E-20 | 2,26 | 3,78 | 3,97E-18 |
| 13353 | 2021 | 33 | 1,92 | 6,93E-05 | 1,37 | 2,72 | 7,69E-04 |
| 40477 | 2021 | 33 | 1,43 | 4,30E-22 | 1,33 | 1,54 | 3,07E-20 |
| 76131 | 2021 | 33 | 1,31 | 1,30E-09 | 1,20 | 1,43 | 3,16E-08 |
| 16321 | 2021 | 33 | 1,20 | 8,18E-05 | 1,09 | 1,31 | 8,85E-04 |
| 40210 | 2021 | 34 | Inf | 2,91E-27 | 23,09 | Inf | 2,56E-25 |
| 45147 | 2021 | 34 | 7,16 | 2,54E-09 | 3,24 | 18,72 | 6,13E-08 |
| 69120 | 2021 | 34 | 3,50 | 1,60E-06 | 1,98 | 6,54 | 2,46E-05 |
| 48143 | 2021 | 34 | 3,17 | 1,46E-08 | 2,05 | 5,03 | 3,18E-07 |
| 81671 | 2021 | 34 | 2,06 | 4,70E-06 | 1,49 | 2,88 | 6,88E-05 |
| 01307 | 2021 | 34 | 1,25 | 2,15E-05 | 1,12 | 1,38 | 2,75E-04 |
| 13353 | 2021 | 35 | 15,69 | 2,76E-04 | 2,44 | 656,11 | 2,64E-03 |
| 92637 | 2021 | 35 | 5,10 | 3,23E-05 | 2,11 | 14,91 | 3,99E-04 |
| 69126 | 2021 | 35 | 4,89 | 7,84E-08 | 2,51 | 10,43 | 1,59E-06 |
| 50935 | 2021 | 35 | 3,55 | 1,18E-40 | 2,89 | 4,39 | 1,68E-38 |
| 66123 | 2021 | 35 | 3,17 | 5,37E-12 | 2,21 | 4,64 | 1,71E-10 |
| 70191 | 2021 | 35 | 2,89 | 8,95E-08 | 1,89 | 4,52 | 1,78E-06 |
| 09221 | 2021 | 35 | 2,04 | 1,84E-04 | 1,38 | 3,06 | 1,84E-03 |
| 40477 | 2021 | 35 | 1,73 | 1,07E-03 | 1,23 | 2,46 | 8,72E-03 |
| 56068 | 2021 | 35 | 1,39 | 1,01E-29 | 1,32 | 1,48 | 1,01E-27 |

Continue on next page.

**Table S4. Continue: showing significant differences in the proportion within the single German postal codes (ZIP) compared between calendar weeks (CW) for S:L452R.**

| zip | YEAR | CW | fold change | p.value | conf.low | conf.high | p.adj |
|---|---|---|---|---|---|---|---|
| 20251 | 2021 | 36 | 4,11 | 2,38E-04 | 1,77 | 11,06 | 2,33E-03 |
| 01307 | 2021 | 36 | 4,02 | 3,31E-05 | 1,92 | 9,45 | 4,06E-04 |
| 14467 | 2021 | 36 | 3,21 | 9,72E-05 | 1,70 | 6,50 | 1,04E-03 |
| 44879 | 2021 | 36 | 3,16 | 1,53E-12 | 2,23 | 4,54 | 5,33E-11 |
| 04779 | 2021 | 36 | 1,86 | 7,81E-05 | 1,35 | 2,59 | 8,51E-04 |
| 92637 | 2021 | 36 | 1,79 | 2,74E-06 | 1,39 | 2,30 | 4,12E-05 |
| 32545 | 2021 | 36 | 1,49 | 4,43E-16 | 1,35 | 1,64 | 2,01E-14 |
| 20095 | 2021 | 37 | 7,59 | 1,04E-04 | 2,27 | 39,77 | 1,11E-03 |
| 50933 | 2021 | 37 | 6,14 | 1,09E-03 | 1,78 | 32,74 | 8,82E-03 |
| 92637 | 2021 | 37 | 4,26 | 1,96E-12 | 2,70 | 6,94 | 6,69E-11 |
| 69126 | 2021 | 37 | 2,06 | 5,00E-21 | 1,77 | 2,42 | 3,33E-19 |
| 48143 | 2021 | 37 | 1,97 | 6,46E-04 | 1,31 | 3,02 | 5,55E-03 |
| 07973 | 2021 | 37 | 1,95 | 7,66E-05 | 1,39 | 2,78 | 8,37E-04 |
| 78467 | 2021 | 37 | 1,61 | 1,51E-13 | 1,42 | 1,83 | 5,53E-12 |
| 70174 | 2021 | 37 | 1,60 | 3,14E-16 | 1,42 | 1,79 | 1,47E-14 |
| 32545 | 2021 | 38 | 14,25 | 9,79E-04 | 2,11 | 608,00 | 8,04E-03 |
| 40210 | 2021 | 38 | 2,72 | 1,84E-16 | 2,11 | 3,54 | 8,76E-15 |
| 85764 | 2021 | 38 | 2,14 | 1,45E-06 | 1,55 | 3,00 | 2,27E-05 |
| 20251 | 2021 | 38 | 1,87 | 9,99E-06 | 1,40 | 2,50 | 1,37E-04 |
| 50933 | 2021 | 39 | 23,70 | 5,75E-28 | 9,82 | 74,59 | 5,38E-26 |
| 92637 | 2021 | 39 | 14,45 | 6,98E-10 | 4,58 | 73,18 | 1,80E-08 |
| 56068 | 2021 | 39 | 9,52 | 4,33E-26 | 5,55 | 17,55 | 3,51E-24 |
| 20251 | 2021 | 39 | 7,22 | 1,08E-05 | 2,53 | 28,25 | 1,46E-04 |
| 78467 | 2021 | 39 | 2,28 | 4,54E-10 | 1,74 | 3,02 | 1,20E-08 |
| 32545 | 2021 | 39 | 1,91 | 2,40E-04 | 1,33 | 2,77 | 2,33E-03 |
| 32105 | 2021 | 39 | 1,79 | 2,28E-04 | 1,30 | 2,48 | 2,24E-03 |
| 40477 | 2021 | 39 | 1,18 | 8,17E-04 | 1,07 | 1,30 | 6,80E-03 |
| 80336 | 2021 | 40 | 8,08 | 3,95E-08 | 3,19 | 26,24 | 8,33E-07 |
| 50858 | 2021 | 40 | 3,99 | 1,86E-08 | 2,31 | 7,25 | 4,04E-07 |
| 16321 | 2021 | 40 | 2,16 | 6,28E-04 | 1,35 | 3,52 | 5,42E-03 |
| 01307 | 2021 | 40 | 2,10 | 2,52E-04 | 1,38 | 3,22 | 2,43E-03 |
| 32545 | 2021 | 40 | 1,81 | 7,49E-04 | 1,27 | 2,60 | 6,30E-03 |
| 01127 | 2021 | 40 | 1,33 | 1,02E-05 | 1,17 | 1,52 | 1,39E-04 |
| 76131 | 2021 | 41 | 2,28 | 2,98E-19 | 1,89 | 2,77 | 1,72E-17 |
| 50933 | 2021 | 41 | 1,68 | 4,87E-05 | 1,30 | 2,18 | 5,61E-04 |
| 69120 | 2021 | 41 | 1,58 | 6,78E-05 | 1,25 | 1,99 | 7,56E-04 |
| 85764 | 2021 | 41 | 1,37 | 2,58E-10 | 1,24 | 1,51 | 6,97E-09 |
| 50858 | 2021 | 41 | 1,30 | 3,91E-05 | 1,15 | 1,48 | 4,65E-04 |
| 92637 | 2021 | 42 | 18,32 | 3,53E-05 | 2,89 | 760,71 | 4,29E-04 |
| 20251 | 2021 | 42 | 6,21 | 6,35E-06 | 2,42 | 20,36 | 9,02E-05 |
| 21502 | 2021 | 42 | 4,75 | 4,19E-04 | 1,79 | 15,83 | 3,82E-03 |
| 55131 | 2021 | 42 | 1,95 | 9,97E-08 | 1,51 | 2,54 | 1,93E-06 |
| 78467 | 2021 | 42 | 1,92 | 3,87E-07 | 1,48 | 2,52 | 6,70E-06 |
| 44879 | 2021 | 42 | 1,39 | 3,45E-15 | 1,28 | 1,51 | 1,48E-13 |
| 09221 | 2021 | 42 | 1,30 | 4,39E-06 | 1,16 | 1,45 | 6,45E-05 |
| 80336 | 2021 | 42 | 1,28 | 7,15E-06 | 1,15 | 1,43 | 1,01E-04 |
| 44137 | 2021 | 43 | 26,83 | 7,16E-20 | 8,81 | 133,20 | 4,38E-18 |
| 32545 | 2021 | 43 | 13,38 | 6,81E-04 | 2,06 | 561,91 | 5,81E-03 |
| 76131 | 2021 | 43 | 8,42 | 9,52E-14 | 4,17 | 19,25 | 3,66E-12 |
| 04779 | 2021 | 43 | 7,44 | 7,31E-05 | 2,27 | 38,47 | 8,08E-04 |
| 09221 | 2021 | 43 | 5,58 | 2,52E-04 | 1,93 | 22,06 | 2,43E-03 |
| 85354 | 2021 | 43 | 4,19 | 3,35E-05 | 1,96 | 9,97 | 4,08E-04 |
| 80336 | 2021 | 43 | 2,80 | 3,85E-11 | 2,02 | 3,94 | 1,11E-09 |
| 87435 | 2021 | 43 | 2,13 | 6,72E-05 | 1,44 | 3,18 | 7,55E-04 |
| 01127 | 2021 | 43 | 1,72 | 3,55E-05 | 1,32 | 2,26 | 4,29E-04 |

Continue on next page.

**Table S4. Continue: showing significant differences in the proportion within the single German postal codes (ZIP) compared between calendar weeks (CW) for S:L452R.**

| zip | YEAR | CW | fold change | p.value | conf.low | conf.high | p.adj |
|---|---|---|---|---|---|---|---|
| 21502 | 2021 | 43 | 1,46 | 2,82E-04 | 1,18 | 1,81 | 2,69E-03 |
| 81675 | 2021 | 43 | 1,23 | 2,40E-05 | 1,12 | 1,36 | 3,04E-04 |
| 69126 | 2021 | 44 | 2,89 | 3,02E-72 | 2,55 | 3,28 | 9,06E-70 |
| 40225 | 2021 | 44 | 2,38 | 3,91E-04 | 1,44 | 4,06 | 3,62E-03 |
| 21502 | 2021 | 44 | 2,19 | 1,64E-04 | 1,42 | 3,42 | 1,66E-03 |
| 30159 | 2021 | 44 | 2,14 | 1,79E-04 | 1,41 | 3,28 | 1,80E-03 |
| 14467 | 2021 | 44 | 1,36 | 1,60E-04 | 1,16 | 1,61 | 1,63E-03 |
| 56068 | 2021 | 44 | 1,34 | 8,07E-04 | 1,13 | 1,59 | 6,73E-03 |
| 76131 | 2021 | 44 | 1,32 | 1,13E-14 | 1,23 | 1,42 | 4,70E-13 |
| 32105 | 2021 | 44 | 1,21 | 2,01E-04 | 1,09 | 1,33 | 2,00E-03 |
| 39124 | 2021 | 45 | 30,83 | 2,91E-116 | 18,49 | 55,51 | 1,75E-113 |
| 50858 | 2021 | 45 | 3,48 | 1,55E-10 | 2,27 | 5,48 | 4,30E-09 |
| 69126 | 2021 | 45 | 3,16 | 4,34E-09 | 2,07 | 4,97 | 1,02E-07 |
| 32105 | 2021 | 45 | 2,90 | 5,04E-04 | 1,56 | 5,37 | 4,45E-03 |
| 69120 | 2021 | 45 | 2,85 | 8,34E-10 | 1,98 | 4,18 | 2,12E-08 |
| 92637 | 2021 | 45 | 2,35 | 3,85E-05 | 1,52 | 3,72 | 4,60E-04 |
| 40477 | 2021 | 45 | 1,40 | 1,22E-18 | 1,30 | 1,51 | 6,66E-17 |
| 06120 | 2021 | 45 | 1,38 | 3,01E-05 | 1,18 | 1,62 | 3,74E-04 |
| 81671 | 2021 | 45 | 1,24 | 1,30E-05 | 1,13 | 1,37 | 1,73E-04 |
| 22081 | 2021 | 46 | 7,84 | 3,10E-97 | 6,16 | 10,10 | 1,33E-94 |
| 21502 | 2021 | 46 | 7,76 | 1,05E-03 | 1,86 | 69,12 | 8,59E-03 |
| 92637 | 2021 | 46 | 4,56 | 1,58E-06 | 2,26 | 10,16 | 2,45E-05 |
| 50858 | 2021 | 46 | 3,40 | 4,63E-19 | 2,53 | 4,62 | 2,57E-17 |
| 89073 | 2021 | 46 | 2,96 | 1,83E-07 | 1,90 | 4,74 | 3,43E-06 |
| 20251 | 2021 | 46 | 1,88 | 1,00E-03 | 1,27 | 2,81 | 8,19E-03 |
| 76131 | 2021 | 46 | 1,69 | 5,12E-10 | 1,43 | 2,02 | 1,33E-08 |
| 07973 | 2021 | 46 | 1,56 | 3,34E-04 | 1,22 | 2,02 | 3,14E-03 |
| 16321 | 2021 | 46 | 1,47 | 4,57E-04 | 1,18 | 1,84 | 4,12E-03 |
| 50933 | 2021 | 46 | 1,35 | 2,82E-13 | 1,25 | 1,47 | 1,02E-11 |
| 78224 | 2021 | 46 | 1,22 | 8,54E-09 | 1,14 | 1,30 | 1,98E-07 |
| 40477 | 2021 | 47 | 4,11 | 1,22E-13 | 2,69 | 6,49 | 4,53E-12 |
| 76131 | 2021 | 47 | 2,28 | 8,64E-08 | 1,65 | 3,18 | 1,73E-06 |
| 50933 | 2021 | 47 | 2,27 | 8,69E-09 | 1,69 | 3,08 | 2,00E-07 |
| 30159 | 2021 | 47 | 1,91 | 2,18E-35 | 1,72 | 2,12 | 2,62E-33 |
| 85764 | 2021 | 47 | 1,77 | 1,02E-07 | 1,42 | 2,21 | 1,94E-06 |
| 04779 | 2021 | 47 | 1,45 | 2,03E-04 | 1,19 | 1,78 | 2,02E-03 |
| 56068 | 2021 | 47 | 1,42 | 9,56E-09 | 1,25 | 1,60 | 2,17E-07 |
| 85354 | 2021 | 47 | 1,24 | 9,45E-08 | 1,14 | 1,34 | 1,85E-06 |
| 06120 | 2021 | 47 | 1,18 | 3,47E-04 | 1,08 | 1,29 | 3,26E-03 |
| 30159 | 2021 | 48 | 5,50 | 5,06E-75 | 4,44 | 6,88 | 1,69E-72 |
| 93053 | 2021 | 48 | 3,23 | 9,36E-07 | 1,93 | 5,64 | 1,52E-05 |
| 69126 | 2021 | 48 | 2,60 | 6,88E-12 | 1,94 | 3,51 | 2,17E-10 |
| 69126 | 2021 | 48 | 2,15 | 1,20E-12 | 1,72 | 2,69 | 4,23E-11 |
| 21502 | 2021 | 48 | 1,65 | 4,70E-04 | 1,23 | 2,23 | 4,22E-03 |
| 40210 | 2021 | 48 | 1,58 | 2,53E-34 | 1,46 | 1,70 | 2,92E-32 |
| 22081 | 2021 | 48 | 1,36 | 8,00E-04 | 1,13 | 1,63 | 6,70E-03 |
| 01307 | 2021 | 48 | 1,19 | 2,27E-05 | 1,10 | 1,30 | 2,90E-04 |
| 40477 | 2021 | 49 | 105,78 | 3,24E-31 | 18,60 | 4118,58 | 3,47E-29 |
| 85354 | 2021 | 49 | 10,64 | 1,86E-07 | 3,35 | 54,23 | 3,46E-06 |
| 76131 | 2021 | 49 | 2,62 | 1,63E-11 | 1,94 | 3,57 | 4,90E-10 |
| 20251 | 2021 | 49 | 2,02 | 4,77E-04 | 1,33 | 3,10 | 4,23E-03 |
| 87435 | 2021 | 49 | 1,96 | 1,94E-06 | 1,47 | 2,64 | 2,97E-05 |
| 39104 | 2021 | 49 | 1,59 | 1,37E-49 | 1,50 | 1,70 | 2,73E-47 |
| 04779 | 2021 | 49 | 1,40 | 3,77E-05 | 1,19 | 1,66 | 4,52E-04 |
| 79104 | 2021 | 49 | 1,26 | 9,31E-11 | 1,17 | 1,35 | 2,63E-09 |

Continue on next page.

**Table S4. Continue: showing significant differences in the proportion within the single German postal codes (ZIP) compared between calendar weeks (CW) for S:L452R.**

| zip | YEAR | CW | fold change | p.value | conf.low | conf.high | p.adj |
|---|---|---|---|---|---|---|---|
| 01458 | 2021 | 49 | 1,17 | 1,50E-05 | 1,09 | 1,26 | 1,97E-04 |
| 80336 | 2021 | 50 | 14,47 | 4,38E-12 | 5,57 | 47,65 | 1,43E-10 |
| 30159 | 2021 | 50 | 2,45 | 2,34E-170 | 2,30 | 2,62 | 2,33E-167 |
| 01458 | 2021 | 50 | 2,02 | 3,32E-05 | 1,42 | 2,89 | 4,06E-04 |
| 44879 | 2021 | 51 | 120,48 | 3,05E-29 | 20,92 | 4697,77 | 2,95E-27 |
| 39104 | 2021 | 51 | 8,98 | 1,96E-46 | 6,24 | 13,24 | 3,46E-44 |
| 76131 | 2021 | 51 | 2,98 | 1,15E-09 | 2,06 | 4,35 | 2,87E-08 |
| 69120 | 2021 | 51 | 1,83 | 1,97E-14 | 1,57 | 2,14 | 8,08E-13 |
| 13353 | 2021 | 51 | 1,70 | 2,98E-05 | 1,32 | 2,21 | 3,72E-04 |
| 04779 | 2021 | 51 | 1,55 | 9,41E-07 | 1,30 | 1,85 | 1,52E-05 |
| 69120 | 2021 | 52 | 2,09 | 4,24E-04 | 1,37 | 3,19 | 3,85E-03 |
| 40477 | 2021 | 52 | 1,96 | 4,21E-04 | 1,33 | 2,89 | 3,83E-03 |
| 76131 | 2021 | 52 | 1,91 | 4,90E-21 | 1,67 | 2,19 | 3,33E-19 |
| 01307 | 2021 | 52 | 1,64 | 1,05E-05 | 1,31 | 2,04 | 1,42E-04 |
| 04779 | 2021 | 52 | 1,48 | 1,40E-05 | 1,24 | 1,78 | 1,86E-04 |
| 40225 | 2021 | 52 | 1,36 | 3,28E-05 | 1,17 | 1,57 | 4,04E-04 |
